# Temporal disparity of action potentials triggered in axon initial segments and distal axons in the neocortex

**DOI:** 10.1101/2022.08.09.503303

**Authors:** Márton Rózsa, Martin Tóth, Gáspár Oláh, Judith Baka, Rajmund Lákovics, Pál Barzó, Gábor Tamás

## Abstract

Neural population activity determines the timing of synaptic inputs, which arrive to dendrites, cell bodies and axon initial segments (AISs) of cortical neurons. Action potential initiation in the AIS (AIS-APs) is driven by input integration, and the phase preference of AIS-APs during network oscillations is characteristic to cell classes. Distal regions of cortical axons do not receive synaptic inputs, yet experimental induction protocols can trigger retroaxonal action potentials (RA-APs) in axons distal from the soma. We report spontaneously occurring RA-APs in human and rodent cortical interneurons that appear uncorrelated to inputs and population activity. Network linked triggering of AIS-APs versus input independent timing of RA-APs of the same interneurons result in disparate temporal contribution of a single cell to in vivo network operation through perisomatic and distal axonal firing.

**One-Sentence Summary:** Network linked triggering of AIS-APs versus input independent timing of RA-APs of the same interneurons result in disparate temporal contribution of a single cell to in vivo network operation.

---

Population signals in neural circuits are predominantly shaped by spatiotemporally concerted action potentials of individual neurons (*1*). Cell class specific action potential timing is characteristic to network states and, conversely, spike timing during network oscillations is used for cell type identification (*2, 3*). Temporally organized synaptic inputs are selectively placed on somatodendritic compartments and on the axon initial segment (AIS) and inputs are considered essential in producing firing in the axon of the postsynaptic cell, in line with the law of dynamic polarization by Ramon y Cajal (*4*). Landmark studies suggest a predominant role for the AIS in input driven spike initiation and action potential dependent output (*5, 6*), however, distal regions of the axon were also suggested to participate in action potential initiation (*7*–*12*). Interneurons of the cerebral cortex do not receive synaptic inputs on axons (*13*) thus local axonal mechanisms were found to be involved in generating retroaxonal action potentials (RA-APs) in the distal axon of GABAergic cells (*11, 12, 14*–*21*). We tested whether compartmentalized mechanisms leading to AIS initiated action potentials (AIS-APs) versus distal axon generated RA-APs result in disparate temporal contribution of a single cell to in vitro and vivo network operation.

Following pioneering work by Sheffield and colleagues in rodents (*17*), we set out to study retroaxonal firing (RAF) in interneurons of the human cortex (*20*). We performed somatic whole-cell patch-clamp recordings of layer 1 interneurons in human cortical brain slices and induced retroaxonal firing (RAF) by injecting repetitive suprathreshold current steps until RA-APs emerged (*17, 19, 22, 23*). We identified two qualitatively different forms of RAF distinguished by the temporal pattern of RA-APs, categorized as persistent or sporadic RAF (Fig. 1, A and B). Similar to previous reports (*17*–*20, 23*), persistent RAF showed barrages of 56 ± 416 RA-APs with high frequencies (13.9 ± 17.3 Hz), however, only a few (8 ± 6) individual RA-APs emerged at low frequencies (0.31 ± 1.5 Hz) during sporadic RAF (Fig. 1, A and B). RA-APs during persistent RAF had lower somatic action potential threshold and lower subthreshold rate of depolarization values compared to AIS-APs (*17*–*20, 23*) We confirmed this during sporadic RAF (Fig. 1C) allowing the separation of RA-APs and AIS-APs throughout this study. Fourth of the tested interneurons (87 of 385 cells, Fig. 1D) in layer 1 of the human neocortex showed RAF, in line with recent experiments (*20*). Morphological recovery permitted post hoc anatomical classification of RAF positive (n=30) and negative cells (n=104) based on light microscopic assessment of biocytin filled cells and indicated that RAF is predominant, but not restricted to human neurogliaform cells (NGFCs). We could trigger RAF in 21 out of 44 NGFCs and 9 out of 90 non-NGFCs (Fig. 1F), and also found that RAF could not be evoked in rosehip cells (n=15) (*20, 24*). We repeated these experiments in the rat neocortex and confirmed the presence of both sporadic and persistent RAF in interneurons predominantly identified as NGFCs (Fig. 1F and fig. S1). We made attempts to localize the site of action potential initiation in anatomically identified human NGFCs (n=2) with simultaneous dual whole cell recordings on the soma and distal axonal bleb of the same cell (Fig. 1, H to K). In response to somatic depolarizing current pulses, AIS-APs were detected first on the somatic electrode, then at the axonal recording site, and the order was reversed for RA-APs (Fig. 1, I to K), and this was accompanied by somatic voltage derivative curves, threshold potentials and rates of depolarization characteristic to AIS-APs and RA-APs (Fig. 1, J and K). Thus, direct measurements confirmed that RA-APs were indeed initiated in the distal axon of human interneurons.

**Fig. 1.**
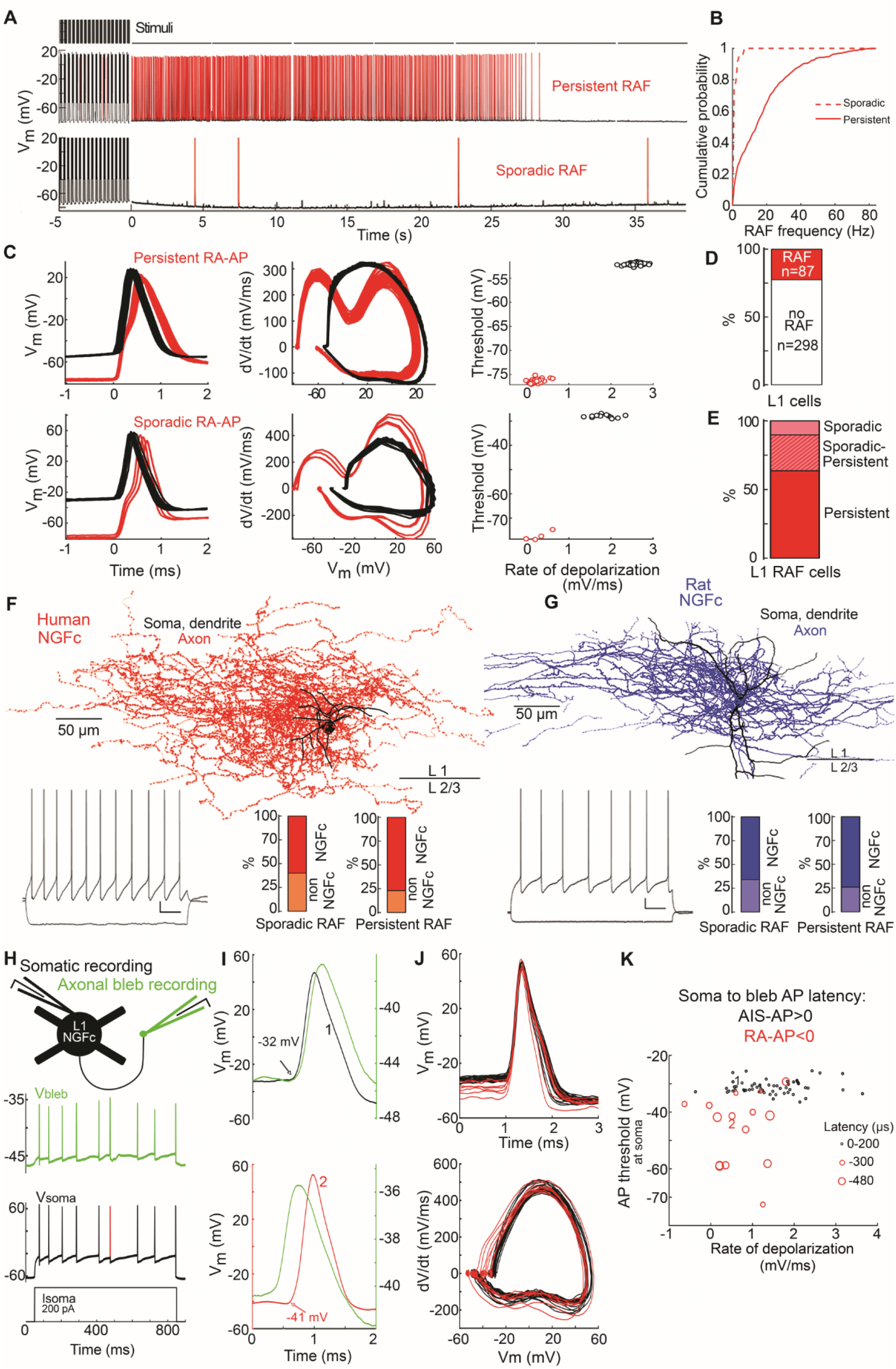
Multiple retroaxonal firing patterns in the human and rat neocortex. (**A**) Human layer 1 interneurons show persistent RAF (middle) and sporadic RAF (bottom) after repetitive suprathreshold somatic current injections (top). (**B**) Cumulative probability of persistent and sporadic RAF frequency shown on panel (A). (**C**) AIS-APs and RA-APs shown on panel (A) are easily distinguishable and show the same kinetics across different types of RAF. Left, Superimposed AIS-APs (black) and RA-APs (red) are aligned according to action potential threshold, respectively. Middle, Phase plot (middle) of AIS-APs (black) and RA-APs (red). Right, Threshold potential and slope of membrane potential 0-1 ms before threshold distinguish AIS-APs and RA-APs. (**D**) Proportion of human layer 1 interneurons with and without RAF detected. (**E**) Individual human interneurons showed persistent only, sporadic only or both RAF patterns. (**F**) Firing pattern (left) and anatomical reconstruction (middle) of a human biocytin-filled layer 1 NGFC showing RAF in the human neocortex (black, soma and dendrites; red, axon). Right, RAF was more prevalent in cells showing NGFC axonal morphology (NGFC: n=21, non-NGFC: n=9). **(G**) Same as in (F) but in rat neocortex. (NGFC: n=22, non-NGFC: n=10). (**H** to **K**) Simultaneous somatic and axonal bleb recordings in the same human NGFC. (**H**) Responses to a somatic current injection (bottom) detected on the soma (black) and the axon (green). An action potential (red) was identified as an RA-AP. (**I**) Top, Somatic detection (black 1) usually preceded axonal detection (green) of the same APs. Bottom, Some APs were also observed with the axonally placed electrode first (green) and with the somatic electrode second (red 2). Note the different somatic threshold potentials for soma first versus second APs (−32 vs. -41 mV). Top and bottom panels show consecutive APs in response to the same somatic current injection. (**J**) Somatic threshold potentials (top) and somatic voltage derivative curves (bottom) separate superimposed AIS-APs (black) and RA-APs (red). (**K**) Soma to bleb latency, somatic threshold potential and rate of depolarization before the action potential differentiates AIS-APs and RA-APs.

Persistent RAF requires intense experimental stimulation protocols (*17, 19, 22, 23*) and was suggested as a mechanism for suppressing epileptiform activity (*19*). How RAF can emerge during physiological conditions is not fully understood (*18, 25*), however sporadic RAF reported here might contribute to brain states with moderate population activity provided that axonal input independent mechanisms drive RA-APs across local threshold. Hyperpolarization activated cyclic nucleotide gated (HCN) channels were found instrumental in generating persistent RAF (*23*) and when we blocked HCN channels by bath-application of ZD7288 (30 µM), the duration of RAF (from 47.2 ± 36.1 to 10.1 ± 16 (s) paired sample t-test p=0.008) and the number of RA-APs (from 624 ± 412 to 48 ± 71, paired sample t-test p=0.007) decreased both in human (n=4) and rodent (n=3) layer 1 interneurons (Fig. 2, B and C). Interestingly, functional expression of HCN channels measured somatically as sag fraction (*26*) was an unreliable predictor of RAF (Fig 2, D and E). Similar to human pyramidal cells (*27*), layer 1 human interneurons (n=358, 0.15 ± 0.29, Mann-Whitney test, p<0.001) showed higher sag fractions compared to those in rat (n=254, 0.04 ± 0.4), but paradoxically, human and rat interneurons with RAF showed smaller sag fractions compared to interneurons without RAF (Fig. 2E, human RAF: n=83, 0.11 ± 0.08, no-RAF: n=275, 0.16 ± 0.16 Mann-Whitney test, p<0.001; rat RAF: n=97, 0.02 ± 0.07, no-RAF: n=157, 0.06 ± 0.11, Mann-Whitney test, p<0.001; fig. S2). Previous recordings showed selective HCN channel expression in axons of mouse fast spiking interneurons (*28*), thus a potentially different somatic versus axonal expression of HCN channels might contribute to the absence or presence of RAF. To test this hypothesis, we simultaneously recorded with electrodes placed on the soma and on a distal axonal bleb (106 ± 56 µm Euclidean soma-bleb distance) of layer 1 interneurons (human, n=2 NGFCs and n=3 unidentified; rat, n=3 NGFCs and n=3 unidentified) and found approximately four times higher sag fractions on the axon (somatic: 0.1 ± 0.05 bleb: 0.4 ± 0.14 Two-Sample t-test p<0.001) (Fig. 2, F to H). These results point to the importance of local axonal HCN channel expression in RAF and suggest that exclusively somatic recordings have a limited power in assessing HCN function in neuron types and subcellular compartments. Apart from intrinsic HCN expression in the axon, extrinsic factors such as locally elevated neuronal activity might produce physiological changes in the local extracellular ionic milieu leading to axonal depolarization and RA-AP initiation (*18, 19, 29*). To this end, we injected ACSF with elevated physiological K^+^ concentration (5 mM) (*30*) while monitoring the membrane potential of layer 1 interneurons >40 µm from the injection site, and as a result, induced sporadic RA-APs transiently in human (n=4) and rat (n=7) layer 1 interneurons (Fig. 2, I to L). Taken together, distal axonal initiation of RA-APs might follow multiple scenarios and emerge during relatively quiescent networks states through axonal HCN channels as well as during active periods of the microcircuit when parts of the axon are depolarized by elevated [K^+^]_out_.

**Fig. 2.**
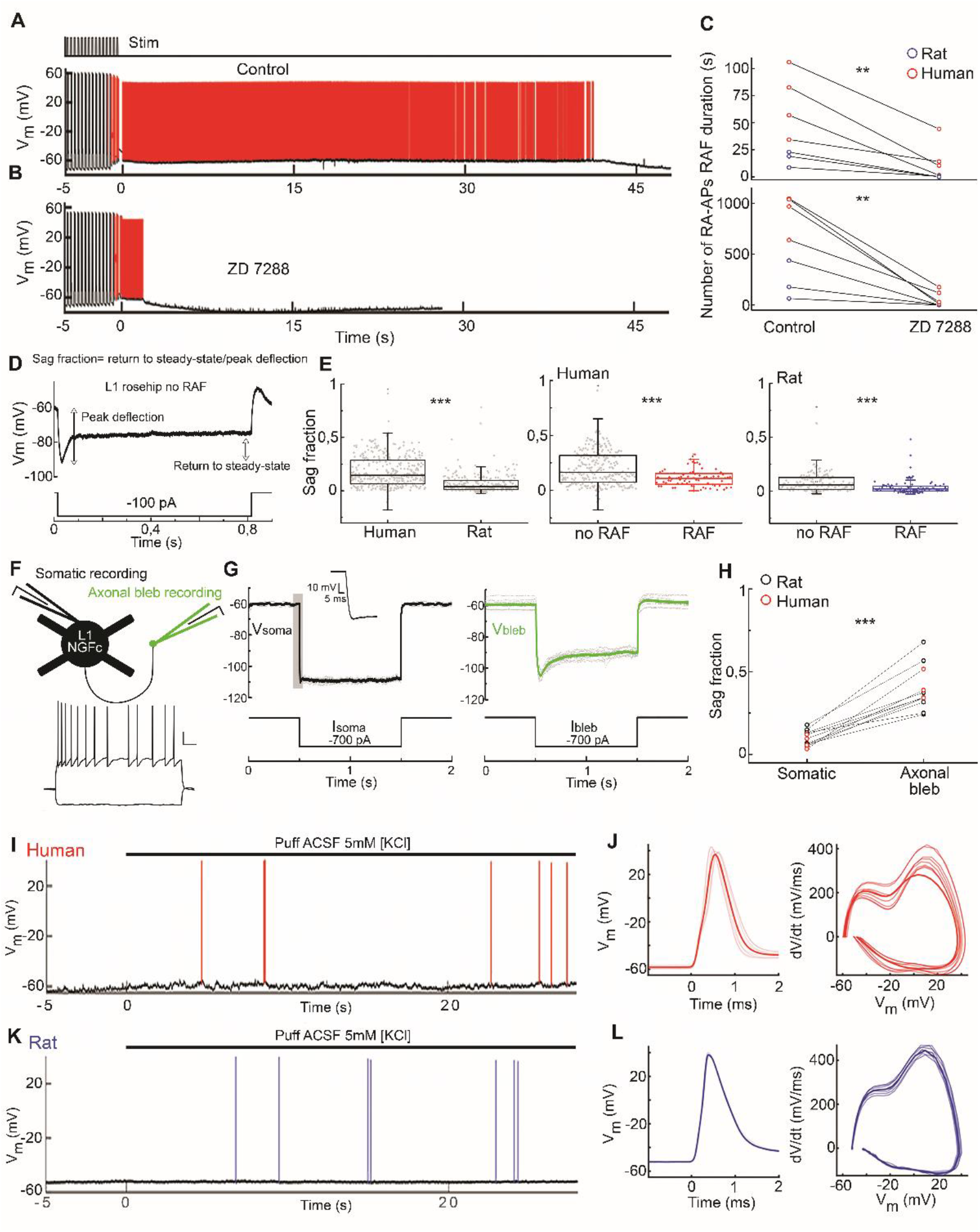
Axonal HCN channels and local increase of potassium ion concentration promote retroaxonal firing. (**A** and **B**) Persistent RAF detected in the human neocortex in control conditions (A) was suppressed by application of the HCN channel blocker ZD7288 30 µM (B). (**C**) The duration of RAF (Paired-Sample t-test p=0.0081) and the number of RA-APs (Paired Sample t-test, p=0.00706) decreased in the presence of ZD7288 relative to control conditions. (**D**) Somatically recorded response of a layer 1 rosehip interneuron to a hyperpolarizing current step with components of the sag fraction indicated. (**E**) Somatic sag fraction was higher in layer 1 interneurons in human compared to the rat (Mann-Whitney test p<0.001). In both species, cells showing RAF had smaller sag fraction compared to cells where RAF could not be induced (Human: Mann-Whitney test p<0.001, Rat: Mann-Whitney test p<0.001). (**F** to **H**) Simultaneous somatic and axonal bleb recordings on the same layer 1 interneurons. (**F**) Experimental design and somatic firing pattern of a human NGFC. (**G**) Representative somatic (black) and axonal (green) responses to somatic and axonal hyperpolarization in the same human NGFC. (**H**) Compared to the soma, the axonal bleb showed higher sag fraction in human and rat cells (Two-Sample t-test, p<0.001). (**I**) Local application of ACSF containing elevated potassium concentration (5 mM KCl) evoked sporadic RAF in somatically recorded human layer 1 interneurons. (**J**) Hyperpolarized threshold potentials and somatic voltage derivative curves identify RA-APs in 5 mM KCl. (**K** and **L)** Same as (I) and (J), but in the rat.

Active brain slice preparations display elevated neuronal activity that might promote physiological RAF and are also regularly used in assessing the contribution of firing in individual neurons to population activity during various network states (*2, 3*). We set out to compare the timing of AIS-APs and RA-APs relative to network behavior using a dual superfusion recording chamber (*31*), in-vivo like [Ca^2+^] and [Mg^2+^] concentrations (*32*) and low cholinergic and dopaminergic tone (see Methods) mimicking neuromodulatory state of deep sleep (*33*) following experiments showing contribution of NGFCs to cortical down states (*34, 35*). In human slices maintained in these conditions, subthreshold membrane potential fluctuations corresponding to cellular up, down and transition states were detected (*36*) in layer 1 interneurons (n=26) recorded somatically without current injections (Fig. 3A). In addition, spontaneous AIS-APS and RA-APs were detected with different threshold potentials and rates of depolarization, respectively (Fig. 3B). Interestingly, AIS-APs and RA-APs were timed differently: AIS-APs were restricted to plateaus of depolarized membrane potential (up states) in all recorded cells with AIS-APs (n=7), however, when occurred (in n=6 cells), spontaneous RA-APs were detected when the membrane potential dropped to hyperpolarized down and transition states in addition to up states. Membrane potential fluctuations were increased 0-50 ms prior to AIS-APs (1.49 mV± 1.76 mV) compared to RA-APs (0.78 mV ± 1.7 mV, Mann-Whitney test p<0.001) presumably due to an increase in synaptic drive during depolarized states (Fig. 3H). In rat slices, somatic recordings in layer 1 interneurons (n=40) were accompanied by layer 5 local field potential (LFP) recordings revealing rhythmic low frequency (1.1 ± 0.55 Hz) LFP deflections synchronized to single cell up states with barrages of synaptic inputs (Fig. 3, F and G). Spontaneous RA-APs were observed in 13 interneurons (all recovered cells were NGFCs, n=5). Similar to experiments in human slices, AIS-APs and RA-APs of rat interneurons were detected with different threshold potentials and rates of depolarization and occurred during up and down states with higher and lower membrane potential fluctuations, respectively (Fig. 3, G to I). Moreover, we found temporal coupling of AIS-APs to near the trough of slow oscillation (average vector length: 0.093, angle: 314°, p<0.001, Rayleigh-test), however, RA-APs showed no phase preference to network oscillations (p=0.07, Rayleigh-test, Fig 3K). Recordings from interneurons (n=90) in slices maintained without external cholinergic and dopaminergic modulators provided similar results (fig. S3), These experiments show that RA-APs occur spontaneously in close to physiological conditions without high intensity stimulation and indicate that AIS-APs and RA-APs of the same interneuron correspond to different temporal domains of ongoing network oscillations.

**Fig. 3.**
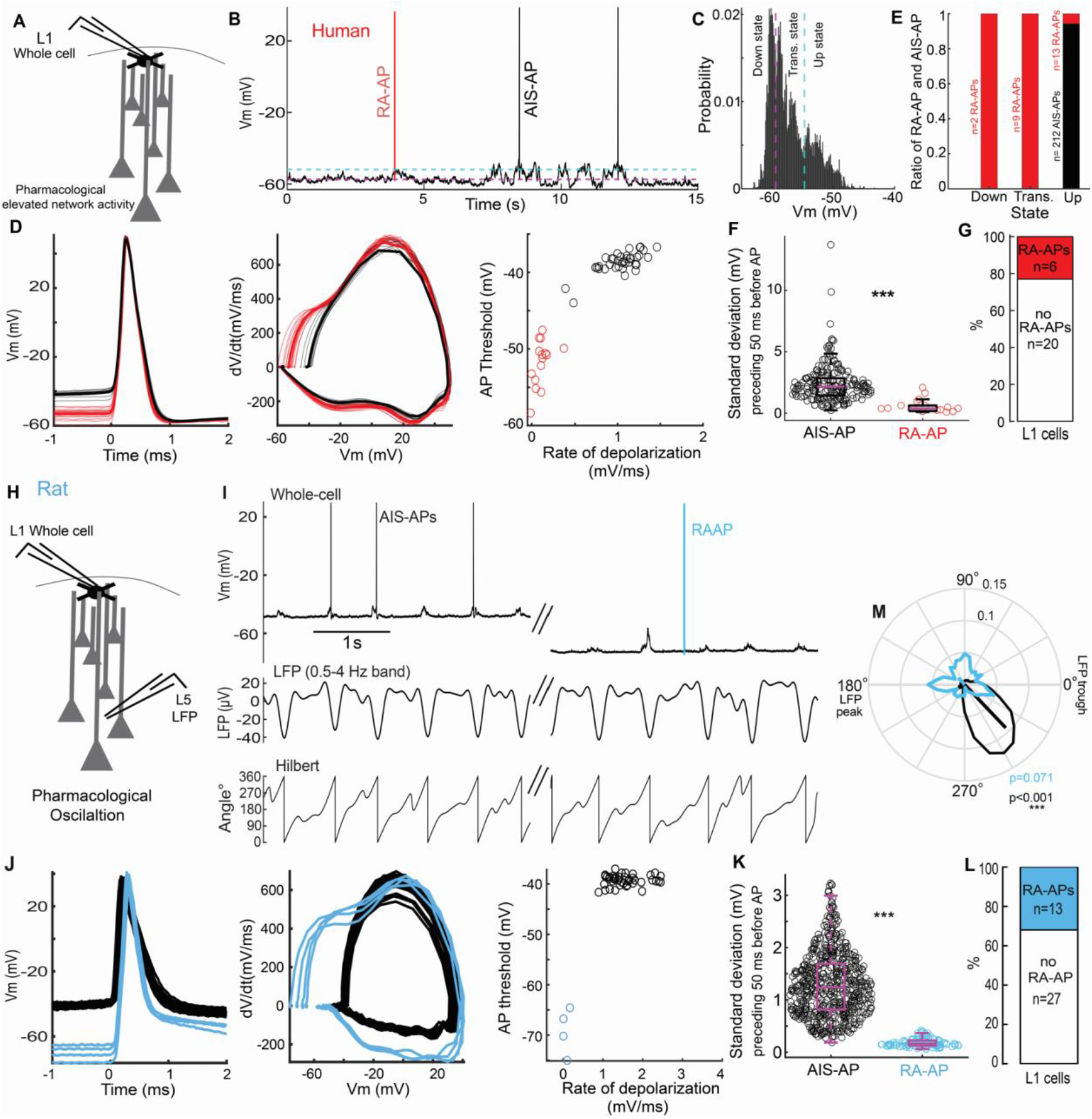
Different temporal domains for AIS-APs and RA-APs in layer 1 interneurons in vitro. (**A** to **G**) Spontaneous AIS-APs and RA-APs detected in active human slices. (**A**) Schematic experimental design. (**B**) AIS-APs (black) and a RA-AP (red) occur at depolarized and hyperpolarized states of membrane potential fluctuations in a human layer 1 interneuron, respectively. Dashed lines separate up, transition and down states. (**C**) Probability of subthreshold membrane potential fluctuations corresponding to cellular up, down and transition states in the human interneuron shown in (B). (**D**) Hyperpolarized threshold potentials (left), different somatic voltage derivative curves (middle) and distinct rates of depolarization (right) identify AIS-APs (black) and RA-APs (red). (**F**) Standard deviation of membrane potential 0-50 ms before APs. (Mann-Whitney test p<0.001). (**G**) Proportion of layer 1 interneurons showing spontaneous RA-APs in active human slices. (**H** to **M**) Detection and timing of AIS-APs and RA-APs in oscillating rat slices. (**H**) Schematic experimental design. (**I**) Simultaneous whole-cell recording from a layer 1 interneuron (top) and LFP recording in layer 5 (middle with Hilbert transformation shown below). A spontaneous RA-AP (blue) occurs during a hyperpolarized state and AIS-APs are synchronized to LFP deflections. (**J**) AIS-APs (black) and RA-APs (blue) were identified according to hyperpolarized threshold potentials (left), different somatic voltage derivative curves (middle) and distinct rates of depolarization (right). (**K**) Standard deviation of membrane potential 0-50 ms before APs (Mann-Whitney test p<0.001). (**L**) Proportion of layer 1 interneurons showing spontaneous RA-APs in active rat slices. (**M**) Circular plot of action potential probability relative to LFP phase (population data: n=22 cells, n=76 RA-APs, n=534 AIS-APs, RA-APs non-uniformity Rayleigh test p=0.071, AIS-APs non-uniformity Rayleigh test p<0.001).

To rule out the possibility that spontaneous RA-APs are brain slice artefacts and to follow up reports that subpopulations of interneurons fire during in vivo down states (*34, 35*), we performed in vivo whole-cell recordings in layer 1 interneurons while simultaneously monitoring layer 5 local field potentials in sleeping and awake mice (Fig. 4A). As expected, interneurons (n=242) recorded in vivo received barrages of excitatory and inhibitory inputs, showed spontaneous membrane potential up and down states and fired most APs at depolarized membrane potentials and during up states (Fig. 4D). However, in 75 (31%) out of the 242 recorded cells, we detected spontaneously occurring APs rising from relatively hyperpolarized membrane potential (Fig. 4D). As above in slice experiments, threshold potentials and rates of depolarization separated two groups of in vivo recorded APs identified as AIS-APs and RA-APs. Overall, layer 1 interneurons fired sporadically (frequency of AIS-APs, 0.65 ± 2.4 Hz; RA-APs, 0.004 ± 0.4 Hz; Mann-Whitney U-test p<0.001) and the proportion of RA-APs represented 3.4% of all in vivo recorded APs in layer 1 interneurons. The ratio of RA-APs varied considerably in individual cells from 0 (n=162 cells) to 100% (n=6 cells) and the average contribution of RA-APs in interneurons with RAF (n=75) was (17 ± 30%). To follow up our slice experiments indicating different temporal distribution for AIS-APs and RA-APs, we tested how these two types of action potentials were related to low frequency (0.5-4 Hz) network activity (Fig. 4, D and J). In line with our findings in vitro, in vivo recorded AIS-APs were coupled to the trough (average vector length=0.019, angle=330°, p<0.001, Rayleigh-test) of population LFP as opposed to RA-APs, that were not significantly coupled to any phase of the slow oscillation (p=0.15, Rayleigh-test, Fig 4J). Thus, RAF occurs spontaneously in vivo in a subpopulation of cortical interneurons. The ratio of AIS-APs and RA-APs swings during slow population oscillations with dominating AIS-APs and RA-APs in up states and down states, respectively.

**Fig. 4.**
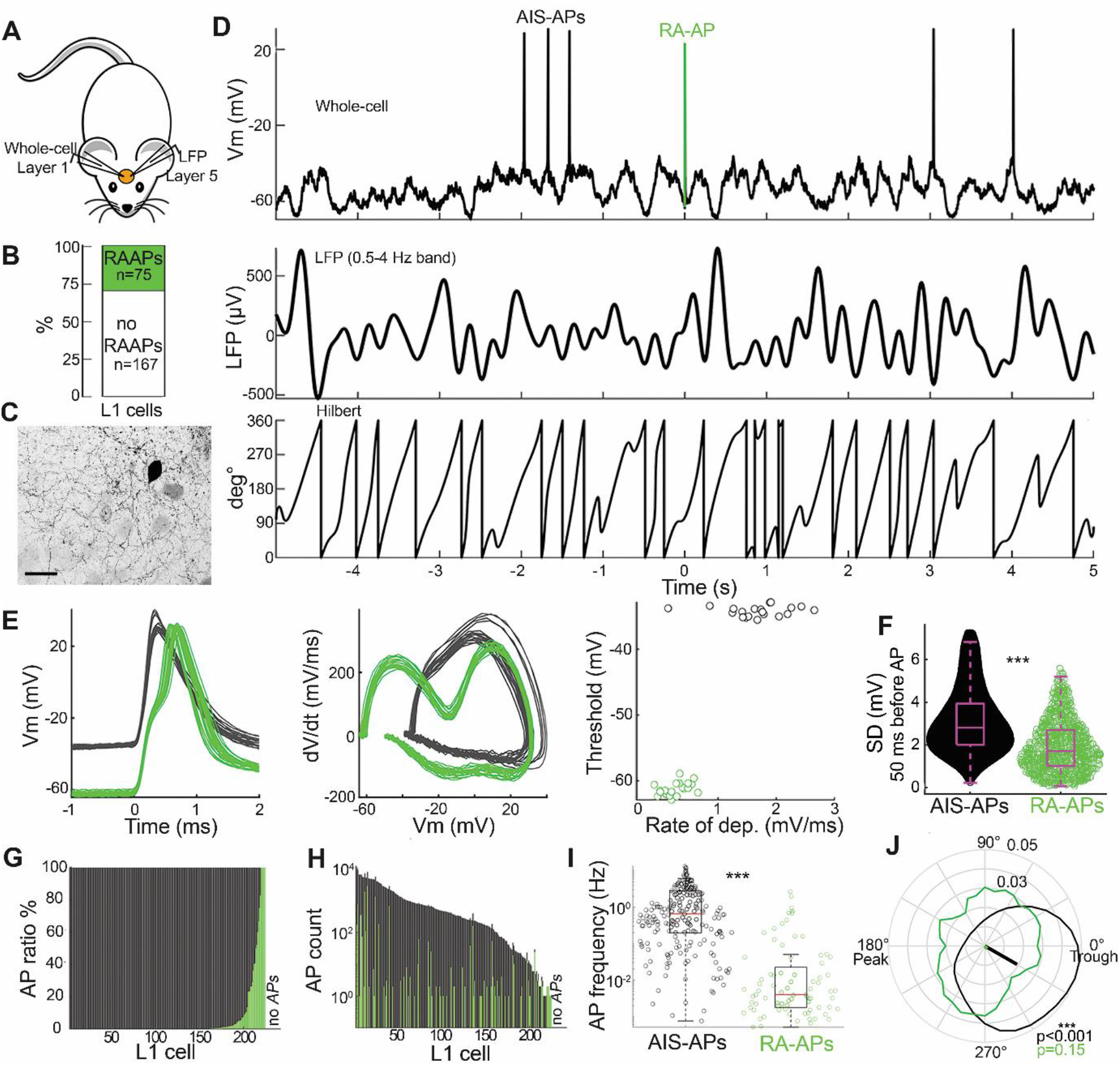
Layer 1 interneurons fire AIS-APs and RA-APs in vivo. (**A**) Schematic of simultaneous layer 1 whole cell and layer 5 LFP recordings in mice. (**B**) Proportion of layer 1 interneurons showing spontaneous RA-APs in the mouse neocortex in vivo. (**C**) Confocal image of a whole-cell recorded layer 1 interneuron that generated RA-APs. (**D**) Simultaneous whole-cell recording from a layer 1 interneuron (top) and LFP recording in layer 5 (middle with Hilbert transformation shown below). A spontaneous RA-AP (green) occurs during a hyperpolarized state and AIS-APs are fired from depolarized up states. (**E**) AIS-APs (black) and RA-APs (green) were identified according to hyperpolarized threshold potentials (left), different somatic voltage derivative curves (middle) and distinct rates of depolarization (right). (**F**) Standard deviation of membrane potential 0-50 ms before APs (Mann-Whitney test p<0.001). (**G** to **I**) Proportion (G), number (H) and frequency (I) of AIS-APs and RA-APs detected in individual layer 1 interneurons. (**J**) Circular plot of action potential probability related to LFP phase (AIS-APs-black, RA-APs-green; non-uniformity Rayleigh test for population data, RA-APs: p=0.15 AIS-APs: p<0.001).

We found that action potentials can be physiologically generated not only in the AIS, but also on distal parts of the axon of cortical layer 1 interneurons. The timing of perisomatic and distal axonal action potential initiation differs. In single cells, membrane potential up state periods are permissive to AIS-APs, but in time windows of cellular down states, suprathreshold activity can be exclusively triggered in the distal axon. Cell to cell jitter of up and down states in individual cells and population level synchronization can result in a rhythmic shift in the balance of AIS-APs versus RA-APs during slow network oscillations. The disparate temporal distribution of AIS-APs and RA-APs enriches the functional repertoire attributable to single cells or cell classes. In line with classic concepts, cell types with AIS-APs selectively contribute to specific phases of ongoing population activity (*2, 3*), whereas cell classes equipped for RA-APs are in position to produce output when somatodendritic input driven AIS-APs cannot reach threshold. Having GABAergic interneurons capable of RA-APs, such mechanisms might contribute to spike triggered GABA release without synaptic inputs otherwise required to drive the cell above threshold. This scenario is likely during cortical down states or during delta waves when most cortical neurons are silent (*37*) and RA-AP dependent GABA release could be helpful in maintaining suppressed population activity. Indeed, neurogliaform cells shown to fire RA-APs (*19*–*21*) were reported to receive thalamic drive at the onset of quiescent down states during slow wave sleep (*34*) and deep cortical NGFCs produced action potentials during down states (*35*). Our results in layer 1 suggest contribution to down state maintenance with RA-AP triggered output (*23, 28*). Furthermore, action potential generation in distal axons is not limited to Down states but might arise in response to direct axonal depolarization by neurotransmitter release (*38*) and, as shown previously (*18, 19, 23, 25*) and here, it can occur during periods of physiologically altered levels of extracellular Ca^2+^ and K^+^ concentrations (*30, 39*). This suggest that distal axons of neurogliaform or other interneurons passing through areas of high K^+^ and low Ca^2+^ extracellular concentrations can generate ectopic action potentials and provide widespread synaptic and non-synaptic GABAergic inhibitory feedback (*40*) to regions of the cortical circuit with elevated activity (*18, 19, 23*).

## Acknowledgments

The authors thank Anna Törteli, Emőke Bakos, Éva Tóth, Katalin Kocsis, Leona Mezei for assistance in anatomical experiments and Attila Ozsvár, Gábor Molnár, Ildikó Szöts, Norbert Mihut, Róbert Averkin, Sándor Bordé for useful feedback and suggestions.

## Funding

Eötvös Loránd Research Network grant ELKH-SZTE Agykérgi Neuronhálózatok Kutatócsoport

Hungarian National Office for Research and Technology grant GINOP 2.3.2-15-2016-00018

Hungarian National Office for Research and Technology grant Élvonal KKP 133807

Hungarian National Office for Research and Technology grant ÚNKP-20-5 - SZTE- 681

National Research, Development and Innovation Office grant OTKA K128863

Hungarian Academy of Sciences, János Bolyai Research Scholarship

National Brain Research Program, Hungary

## Author contribution

Conceptualization: M.R., M.T., and G.T.

Data curation: M.R., M.T., and G.O.

Formal analysis: M.R., M.T., G.O., and G.T.

Funding acquisition: G.T.

Investigation: M.R., M.T., G.O., J.B., and R.L.

Methodology: M.R., M.T., G.O., J.B., and R.L.

Project administration: G.T.

Resources: P.B. and G.T. Supervision: G.T.

Validation: M.R., M.T., and G.T.

Visualization: M.R., M.T., and G.T.

Writing - original draft: M.R., M.T., and G.T.

Writing - review & editing: M.R., M.T., and G.T.

## Competing interests

The authors declare that no competing interests exist.

## Materials and Methods

### Slice preparation

All procedures were performed according to the Declaration of Helsinki with the approval of the University of Szeged Ethical Committee. We used neocortical tissue surgically removed from patients (n =38 female and n =29 male, aged 50 ± 20 years) as part of the treatment protocol for aneurysms, shunt, and brain tumors. Anesthesia was induced with intravenous midazolam and fentanyl (0.03 mg/kg, 1–2 mg/kg, respectively). A bolus dose of propofol (1–2 mg/kg) was administered intravenously. To facilitate endotracheal intubation, the patient received 0.5 mg/kg rocuronium. After 120 s, the trachea was intubated, and the patient was ventilated with a mixture of O_2_ –N_2_O at a ratio of 1:2. Anesthesia was maintained with sevoflurane at monitored anesthesia care volume of 1.2–1.5. Human cortical tissue blocks were removed from frontal (n = 29), temporal (n = 18), occipital (n=3), and parietal (n =4) areas and each sample underwent neuropathological cross examination as part of a dataset published earlier *(26)*. In rodent experiments, we used the neocortical tissue of Wistar rats (postnatal day 18-44, 22 ± 5). Animals were anesthetized by inhalation of halothane, and following decapitation, tissue blocks were prepared from the somatosensory cortex. Tissues were immersed in an ice-cold solution containing (in mM) 130 NaCl, 3.5 KCl, 1 NaH_2_PO_4_, 24 NaHCO_3_, 1 CaCl_2_, 3 MgSO_4_, 10 d(+)-glucose, saturated with 95% O_2_ and 5% CO_2_. Slices were cut perpendicular to cortical layers at a thickness of 350-400 µm with a vibrating blade microtome (Microm HM 650 V) and were incubated at 36 °C for half an hour in the same solution and then for half an hour in a solution composed of (in mM) 130 NaCl, 3.5 KCl, 1 NaH_2_PO_4_, 24 NaHCO_3_, 1 CaCl_2_, 3 MgSO_4_, 10 D(+)-glucose. After the end of incubation, slices were kept at room temperature until use. The solution used during recordings differed only in that it contained 3 mM CaCl_2_ and 1.5 mM MgSO_4_ in the conventional, or 1.2 mM CaCl_2_ and 1 mM MgSO_4_ in the in-vivo like ACSF.

### In vitro electrophysiological recordings

Somatic whole-cell current-clamp recordings were obtained at approximately 36 °C in the conventional sub-merged chamber or dual-super fusion chamber *(31)*. Neurons of layer 1 of cortical slices were targeted, and the recorded cells were visualized by infrared differential interference contrast video microscopy (Olympus BX60WI microscope, Hamamatsu CCD camera equipped with micromanipulators Luigs and Neumann, 652 Ratingen, Germany) at depths of 60–130 µm from the surface of the slice. Micropipettes (3–5 MΩ) were filled with intracellular solution containing (in mM) 126 potassium-gluconate, 4 KCl, 4 ATP-Mg, 0.3 GTP-Na_2_, 10 HEPES, 10 phosphocreatine, and 8 biocytin (pH 7.20; 300 mOsm). Signals were filtered at 10 kHz (Bessel filter), digitized at 50 kHz, and acquired with Patchmaster (HEKA Elektronik GmbH, Lambrecht) software.

### Induction protocol of retroaxonal firing

The induction protocol consisted of 100 ms long repeated square pulses with 300 ms inter-stimulus intervals. Suprathreshold current amplitude was adjusted to evoke at least 4 action potentials during a single pulse (50-500 pA). The induction protocol was terminated, and continuous recording was started upon registering action potentials during the inter-stimulus interval.

### Pharmacology

ZD7288 (HCN channel blocker, 30 μM), carbamylcholine chloride (cholinergic agonist, 2 μM) and SCH23390 (D1 receptor antagonist, 10 μM) were applied in the bath solution. All drugs were obtained from Sigma-Aldrich or Tocris Bioscience.

### Post hoc anatomical analysis

The in vitro electrophysiological recorded cells were filled with biocytin, which allowed post-hoc anatomical analysis after visualizing the cells with DAB reaction. We categorized layer 1 interneurons as neurogliaform cells, non-neurogliaform cells, or rosehip cells (*2*) types. The NGFCs were identified by their relatively small somata, dense and local axonal arborizations, and their thin axons bearing a large number of boutons *(24)*. The non-NGFCs had relatively large somata and sparse axonal arborizations, and their axons usually reached other cortical layers as well. We identified the rosehip interneurons as having large axonal boutons forming very compact, bushy arborizations *(24)*.

### Axonal bleb recordings

Bleb recordings were obtained with the same in vitro slicing and incubation protocol as described above. For visualization of axonal compartments, we performed whole-cell patch clamp configuration (3-5 MOhm pipettes) and filled the layer 1 interneurons with an internal solution containing Alexa Fluor 594 (30 µM). After ∼15 minutes of waiting for dye spreading we targeted blebs of axons under two-photon microscopy (Zeiss Axio Examiner LSM7; Carl Zeiss AG, Oberkochen, Germany, 40× water-immersion objective 1.0 NA; Carl Zeiss AG, Oberkochen, Germany, driven by a Mai Tai DeepSee [Spectra-Physics, Santa Clara, CA] femtosecond Ti:sapphire laser tuned to 800 nm) with 20-25 MOhm micropipettes filled with an internal solution containing Alexa Fluor 488 (20 µM). Next, we injected hyperpolarization current steps alternately to soma and bleb, or injected depolarization current steps only in soma.

### In vitro oscillations

In order to induce slow-wave oscillation in vitro, we used dual-superfusion slice chamber *(31)* and modified the recording solution: 1.2 mM CaCl_2_ and 1 mM MgSO_4_ (*4*). We induced the emergence of oscillation in the delta frequency band (1-4 Hz) with additional cholinergic agonist carbachol (2 µM) and dopamine receptor 1 antagonist SCH23390 (10 µM) *(33)*. We measured local field potential (LFP) in layer 5 with micropipettes (1-2 MΩ) which were filled with CaCl_2_ free recording solution. Signals were filtered at 10 kHz (Bessel filter), digitized at 50 kHz with Patchmaster software (HEKA Elektronik GmbH, Lambrecht). In some cells n=16, steady state somatic current injections were needed to cross AIS-AP threshold, RA-APs were absent during these periods.

### In vivo electrophysiological recordings

All procedures were performed according to the Declaration of Helsinki with the approval of the University of Szeged Ethical Committee. Young adult (postnatal day 35-180, mean: 81±30 days) male and female mice (GAD67-GFP line G42 n=37, Ai9 n=5, C56/B6J n=3, vGAT-AI9 n=75) were anesthetized using isoflurane (2.5% for induction, 1.5% during surgery), and a custom metal head post was cemented to the skull. After full recovery, mice were placed on water restriction and got water only in the recording rig. After a week of habituation, mice were anesthetized using isoflurane, or with a mixture of ketamine-xylazine (0.1 mg ketamine and 0.008 mg xylazine, or with 0.4 mg chloral hydrate per gram body weight), and a circular craniotomy (2-3 mm diameter) was made above the dorsal cortex. We performed a durotomy, filled the craniotomy with 1.5% agarose, then a bar shaped coverslip was secured on top to suppress brain motion and leave access to the brain on the lateral sides of the craniotomy for the intracellular and extracellular electrodes. The craniotomy was covered and mice recovered in their homecage or in the recording rig before awake recordings. For anesthetized recordings, mice were transferred to the recording rig right after surgery while keeping the mouse anesthetized. Awake whole cell recordings were performed at least 1-2 hours after the surgery. Mice were video monitored and not required to do any task, they were sitting in the recording rig comfortably, occasionally grooming, sometimes even falling asleep. LFP recording micropipettes (1-3 MΩ) were filled with calcium-free ACSF. Local field potential recordings were performed in deeper layers of cortex (electrode depths: 548.06 ± 348.26 µm). Intracellular micropipettes (3–6 MΩ) were filled with intracellular solution containing (in mM) 126 potassium-gluconate, 4 KCl, 4 ATP-Mg, 0.3 GTP-Na2, 10 HEPES, 10 phosphocreatine, and 8 biocytin (pH 7.20; 300 mOsm) and with fluorescent dye (10 µM Alexa 594 or Alexa488, depending on the transgenic line used). Somatic whole-cell recordings were obtained from layer 1 interneurons 20-120 μm depth from brain surface, in a ∼1000 micrometer (806.7 ± 506 µm) range of the LFP electrode. Cells were visualized with either two-photon microscopy (Femtonics galvo-galvo two photon microscope built on an Olympus BX61WI upright microscope base, driven by a Mai Tai DeepSee (Spectra-Physics, Santa Clara, CA] femtosecond Ti:sapphire laser at 800-850 nm) or infrared oblique illumination. The infrared LED (Osram SFH 4550, 850 nm) illuminated the craniotomy 30-45° from normal. Warm saline (35-37°C) was perfused in the craniotomy to keep the cortex at physiological temperature. We recorded in current clamp mode, signals were filtered at 10 kHz (Bessel filter), digitized at 50 kHz, and acquired with Patchmaster (HEKA) software.

### Post Hoc anatomical analysis of in vivo recordings

After the whole cell recording experiments, mice were deeply anesthetized with a mixture of ketamine-xylazine (0.1 mg ketamine and 0.008 mg xylazine per gram body weight), and transcardially perfused first with cold 0.1M PB, then with freshly prepared PB containing 4% paraformaldehyde and 0.15% glutaraldehyde. The brain was removed and fixed overnight in the same solution. We sectioned 50 µm thick coronal slices at the craniotomy, then the slices were frozen in liquid nitrogen and thawed in 0.1 M PB. For visualization of the neurobiotin filled cells, we used Streptavidin conjugated Alexa488 or Cy3 (Jackson ImmunoResearch) in 1:400 dilution in TBS overnight, and mounted in Vectashield (Vector Laboratories). Slides were imaged on a Zeiss LSM 880 confocal microscope using 40x oil immersion objective (1.4 NA).

### Data analysis

We analyzed the electrophysiological data with Fitmaster software (HEKA Elektronik GmbH, Lambrecht), and custom MATLAB (The Math Works, Inc.) scripts.

### Analysis of retroaxonal firing patterns

Action potentials were detected on unfiltered whole-cell traces. We extracted threshold potential at the onset of APs when the depolarization rate exceeded 5 mV/ms and calculated the rate of depolarization as the slope of a linear fit of the trace in the last 1 ms before the threshold. The threshold potential and rate of depolarization distinguishes between AIS-APs and RA-APs, and we could confirm the separation of AIS-APs and RA-APs by looking at the first order derivative of the APs. Successful RAF inductions were categorized based on the firing frequency of RA-APs, with a threshold of 5 Hz. After categorization into two groups (sporadic, <5 Hz, tonic, >5 Hz) we extracted the following parameters for all RAF epochs: number of required action potentials for the induction of RAF (number of evoked AIS-APs during stimulation), number of RA-APs (number of RA-APs during continuous recording), RAF duration (elapsed time between the first and the last RA-APs during continuous recording), RAF frequency (number of RA-APs divided by the RAF duration), maximal instantaneous firing frequency (the maximum of the reciprocal of the inter-spike intervals during RAF epoch).

### Analysis of AP timing related to network oscillations

Action potentials recorded during network oscillations in vivo and in vitro were categorized as *RA-APs* and *AIS-APs* as above. Next, we generated 4 seconds long action potential triggered LFP traces around each AP, calculated their power spectrum using fast Fourier transform and corrected the power values with the frequency values to compensate for the 1/f nature of the spectral power. For calculating slow oscillation phase coupling, we considered only action potentials where the corrected power ratio of the delta band (0.5-4 Hz) was bigger, than 0.1, compared to the full range of frequencies (0.5-45 Hz) *(41)*. Selected raw LFP traces were bandpass filtered with Butterworth filter in the delta frequency range (0.5-4 Hz) and partitioned in 0-360° range with Hilbert transformation. The LFP troughs denote the UP-state (phase: 0° and 360°), and the LFP peak denotes the DOWN-state (phase: 180°) throughout the manuscript. We used the phase at the peak of each action potential for further analysis. We calculated the mean vector direction and length with MATLAB Circular Statistics Toolbox scripts *(42)* to extract phase preferences of AIS-APs and RA-APs. Phase distribution plots were binned with a 10° window and smoothed with a moving average with a 30° window. We determined non-uniformity of the *APs* phase coupling with Rayleigh-test *(42)*.

### Statistics

We presented the data as median ± standard deviation unless otherwise stated. All statistical analyses were performed with MATLAB Statistics and Machine Learning Toolboxes. We stated statistical difference if p<0.05 * denotes p<0.05, ** denotes p<0.01, *** denotes p<0.001 throughout the figures. For all statistical tests, we first tested for normality with the Kolmogorov-Smirnov test. If data was judged to be normally distributed, we performed paired t-test or two-sample t-test, otherwise, we used the non-parametric Mann–Whitney-U test. Phase coupling of action potentials was tested with MATLAB circular statistics toolbox *(42)*. We applied the Rayleigh test to identify the possible phase coupling of action potentials and the non-uniformity.

**Fig. S1.**
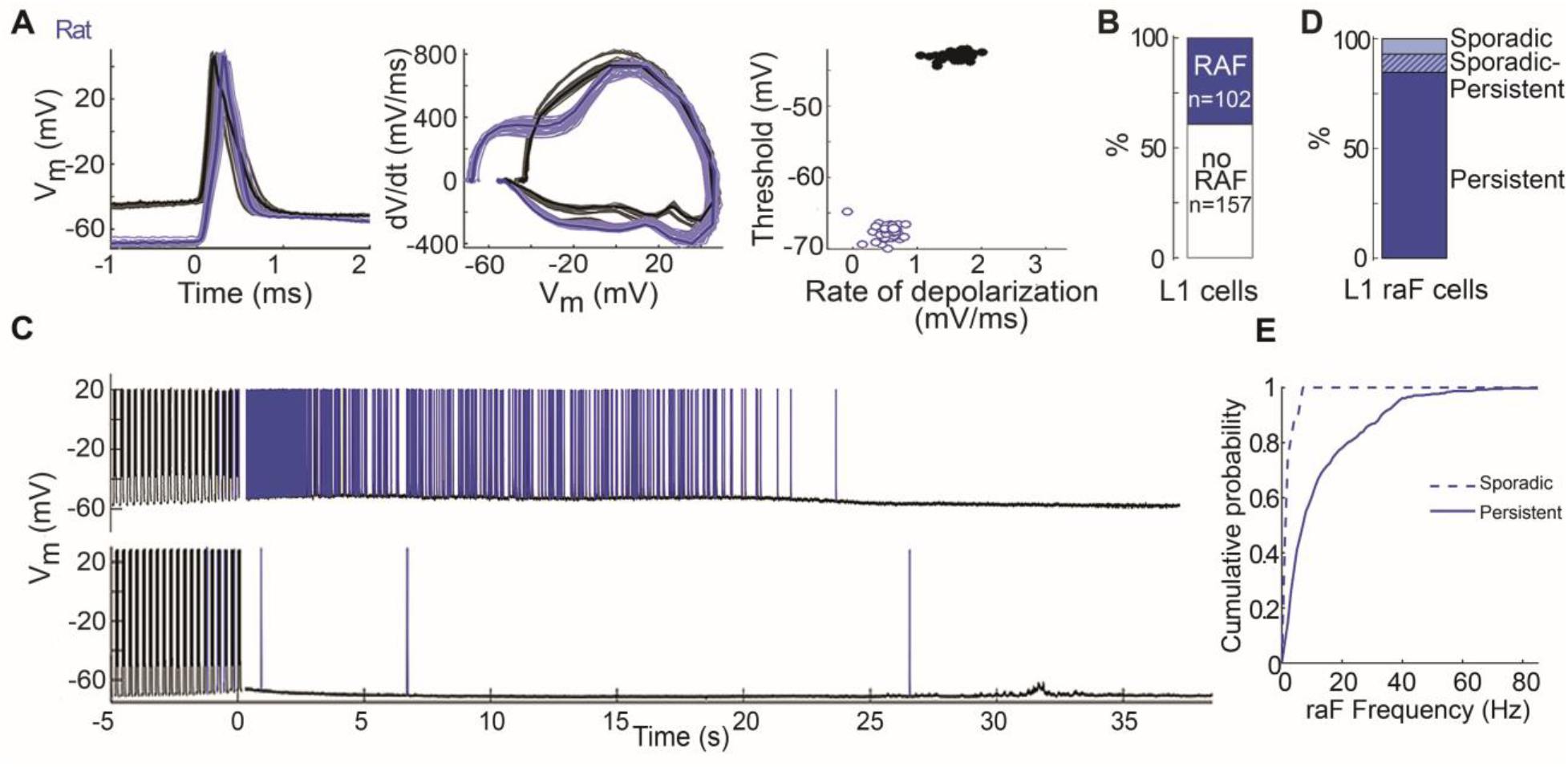
Multiple RAF patterns in the rat neocortex. **(A)** AIS-APs and RA-APs are easily distinguishable and show the same kinetics across different types of RAF in layer 1 interneurons of the rat cortex. Superimposed AIS-APs (black) and RA-APs (blue) of a representative cell are aligned to action potential threshold (left). Phase plot (middle) of AIS-APs (black) and RA-APs (blue). The threshold potential and slope of voltage trace in the last 1 ms before threshold separate AIS-APs and RA-APs. **(B)** Approximately a third of the tested cells (102 out of 259) showed RAF. (**C**) We observed both sporadic RAF (RAF frequency: 1 ± 1.8 Hz, Number of RA-AP: 7± 2.2) and tonic RAF (RAF Frequency: 6.9 ± 14 Hz, Number of RA-AP: 100 ± 241) in the rodent brain slice preparation. Two representative layer 1 interneurons show different RAF after repetitive suprathreshold somatic current injection: tonic RAF (top) and sporadic RAF (bottom). (**D**) Percentages of RAF interneurons showing sporadic, tonic RAF, or both. (**E**) Cumulative probability for tonic and sporadic RAF frequency.

**Fig. S2.**
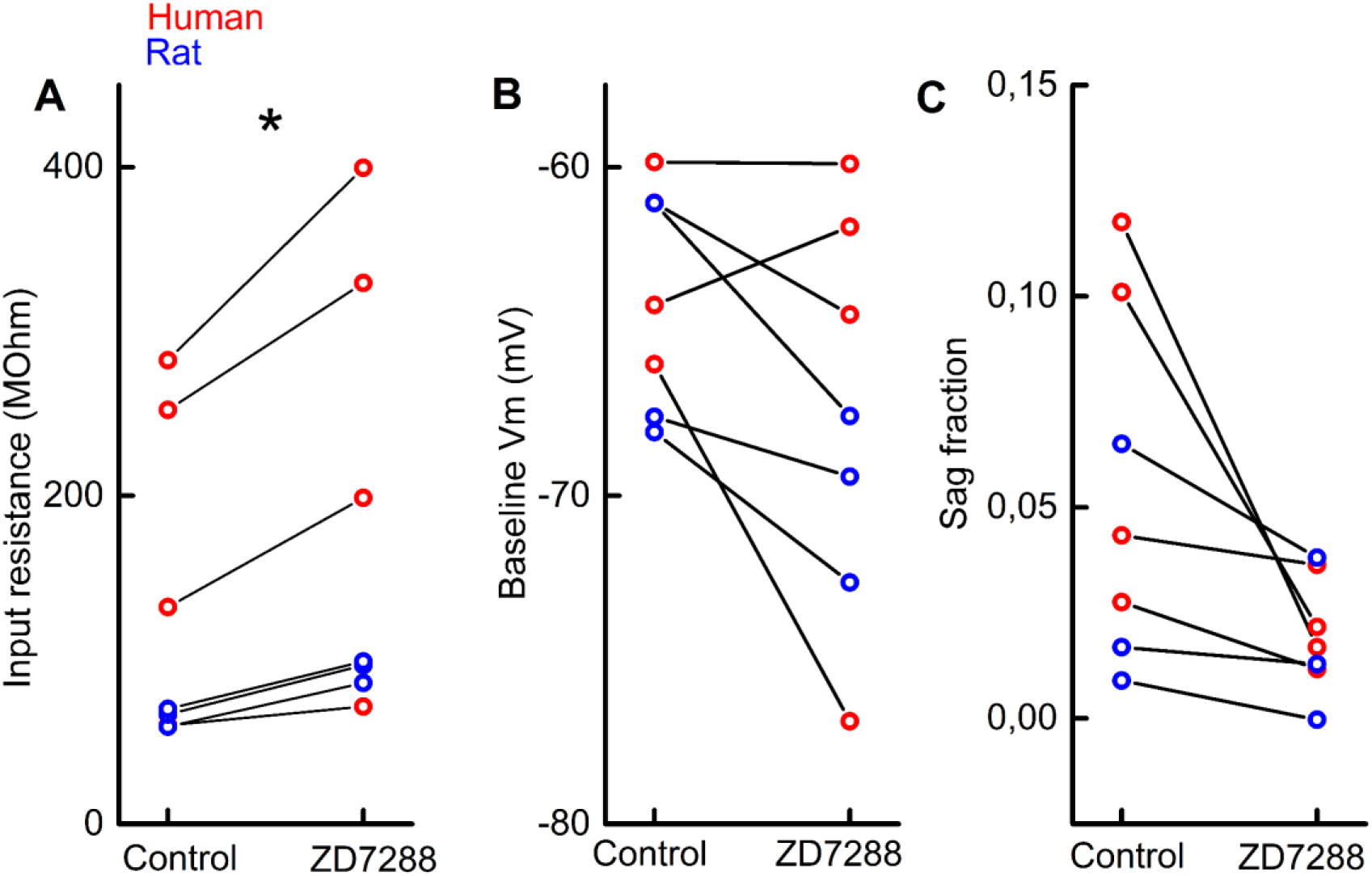
Effect of HCN channel blocker ZD7288 on electrophysiological properties of layer 1 interneurons in the human and rat neocortex. Application of ZD7288 (30 µM) slightly but significantly increased the input resistance and slightly hyperpolarized their resting membrane potentials and decreased sag fraction of the recorded neurons. (**A**) Input resistance (control: 70 ± 96 MOhm, ZD7288: 98 ± 132 MOhm, paired-sample t-test p=0.01). (**B**) Resting membrane potential (control: -60 ± 3 mV, ZD7288: -67 ± 6 mV paired sample t-test p=0.075). (**C**) Sag fraction (control:0.04 ± 0.04, ZD7288:0.017 ± 0.014 paired sample t-test p=0.057).

**Fig. S3.**
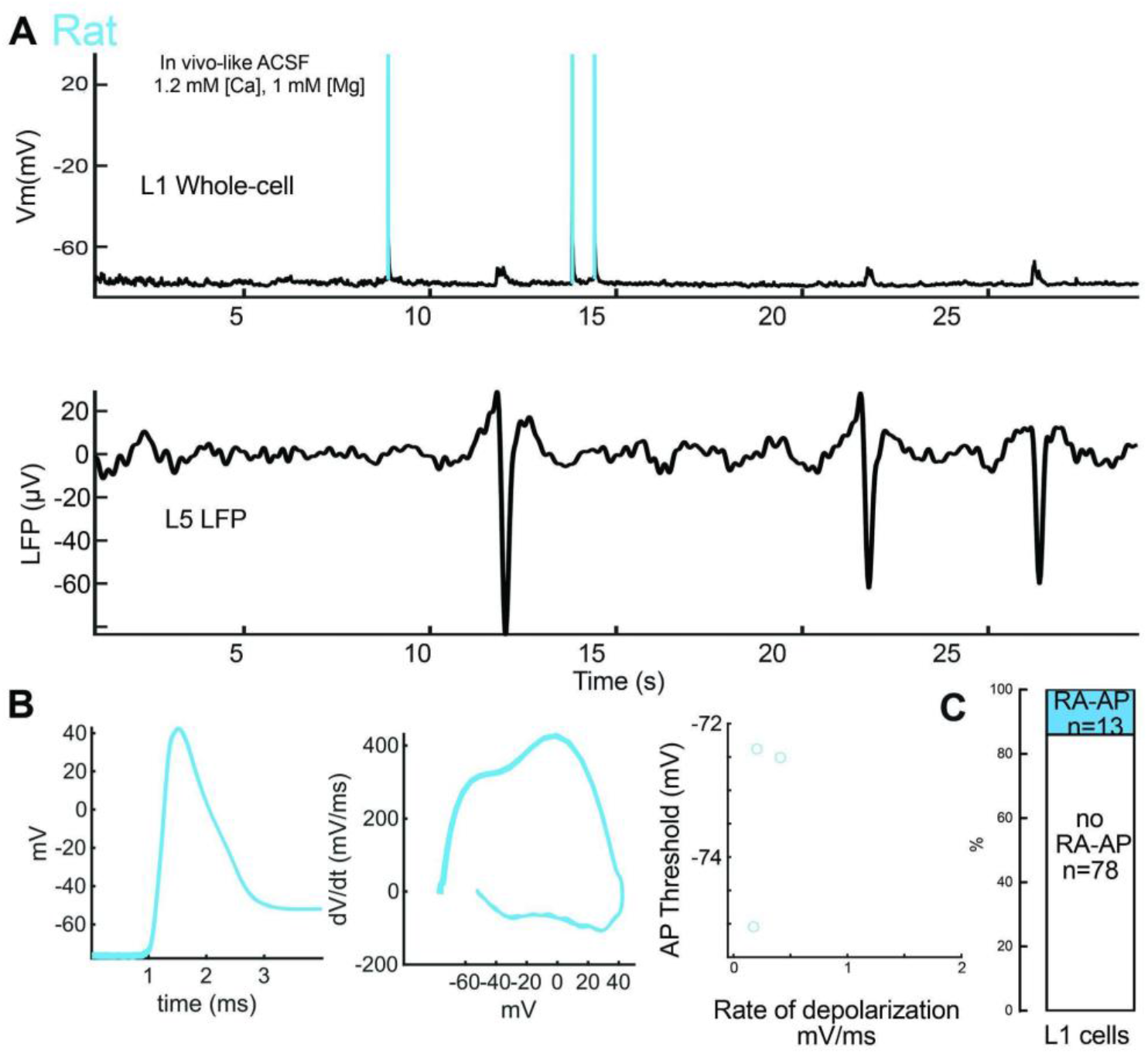
Elevated network activity can be evoked without pharmacological interventions in the rat neocortex. In vivo-like ACSF and dual-superfusion chamber provides suitable environment for the elevation of network activity and the appearance of spontaneous RA-APs, without the addition of pharmacological agents. (**A**) Layer 1 interneuron in the rat neocortex fires RA-APs during elevated network activity. Bottom shows LFP trace measured in layer 5. Low frequency (0.11 ± 0.04 Hz) Up-states are visible on the LFP recording as negative deflections, and as subthreshold peaks in the intracellular recording (top). (**B**) RA-APs (cyan) aligned to threshold (left), and phase plot shows characteristic kinetics (right). Threshold potential and rate of depolarization values are characteristic to RA-APs (bottom). **(C)** Proportion of layer 1 interneurons showing spontaneous RAAP. RA-AP cells: n=4 NGFc, n=5 non-NGFc.

**Fig. S4.**
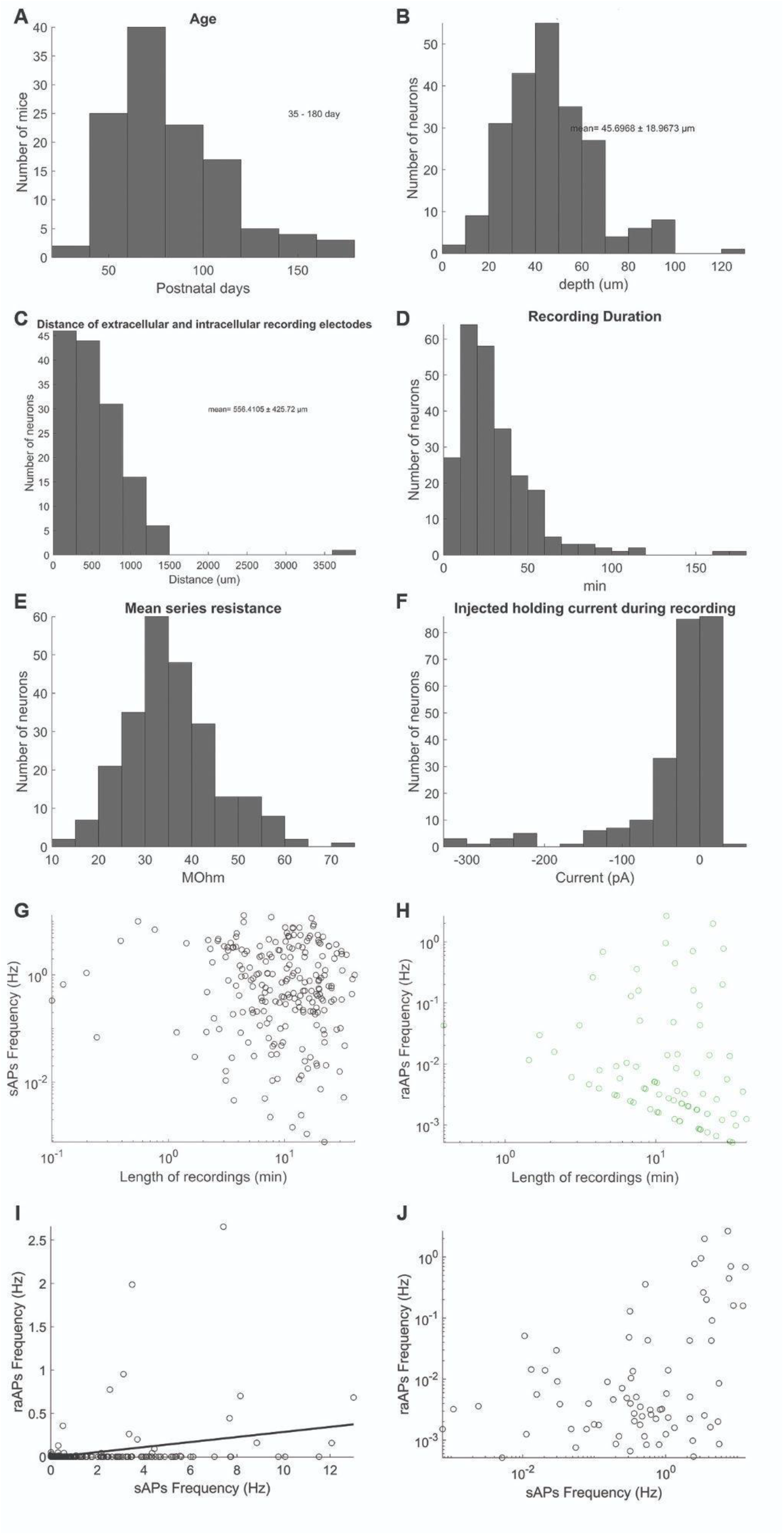
Statistics of in vivo recordings. (**A**) Age of animals (postnatal days). (**B**) Depth of whole-cell recorded layer 1 interneurons from the surface of the brain. (**C**) Distance between LFP and intracellular recording electrodes from each other. (**D**) Duration of whole cell recordings. (**E**) Mean series resistance during recordings. **F:** Injected holding currents during recordings. (**G**) AIS-APs frequency vs the duration of the recording. (**H**) RA-APs frequency vs the duration of the recording. (**I**) Frequency of AIS-APs and RA-APs (R=0.29, p<0.001) (**J**) Frequency of AIS-APs and RA-APs on a logarithmic scale.

**Fig. S5.**
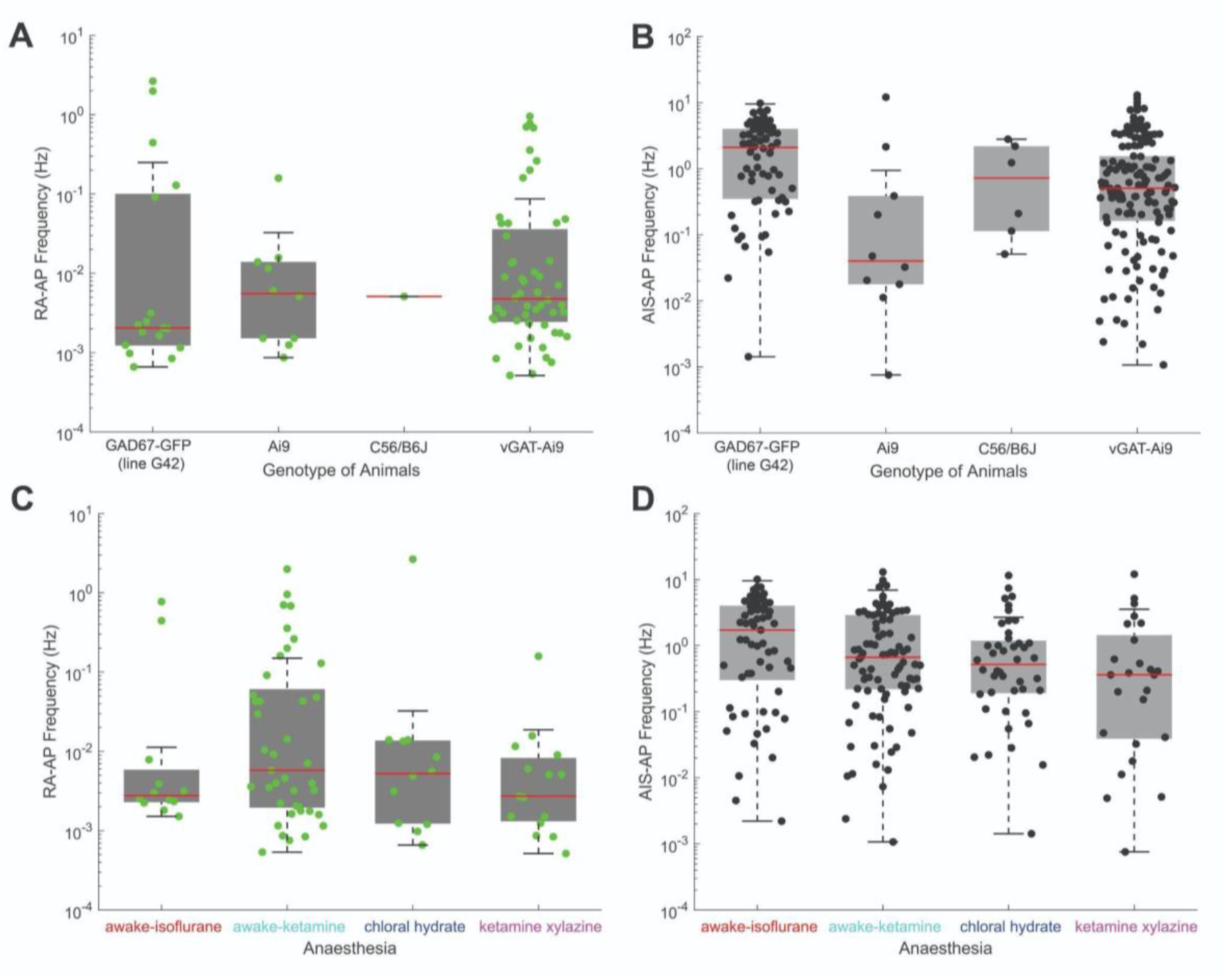
RA-AP occurrence cannot be explained by genotype or anesthesia. (**A** and **B**) RA-AP and AIS-AP frequencies for each genotype tested, each dot represents a neuron. (**C** and **D**) RA-AP and AIS-AP frequencies for each condition tested: awake recordings after isoflurane or ketamine-xylazine anesthesia, or recordings during chloral hydrate or ketamine-xylazine anesthesia, each dot represents a neuron.

## Notes

### Competing Interest Statement

The authors have declared no competing interest.

